# A simple method for introducing a cutoff to hydrodynamic interactions in Brownian dynamics simulations

**DOI:** 10.1101/2025.05.31.657166

**Authors:** Adrian H. Elcock

## Abstract

Brownian dynamics (BD) simulations that include hydrodynamic interactions (HIs) modeled at the Rotne-Prager-Yamakawa (RPY) level of theory are a valuable tool for accurately modeling the translational and rotational diffusion of macromolecules such as proteins and nucleic acids. A major drawback to the inclusion of HIs in BD simulations is their computational expense, and an obvious way to consider reducing the expense of BD-HI simulations is to include a cutoff such that HIs beyond a certain distance are omitted. Unfortunately, a naïve attempt to implement such a scheme usually leads to the RPY diffusion tensor becoming non-positive definite, which has the consequence that it becomes impossible to compute the correlated random displacements required by the Ermak-McCammon BD-HI algorithm. Here I show that a simple approach can be used to overcome this problem and implement a distance-based cutoff scheme that is guaranteed to lead to a diffusion tensor that is positive definite. The method involves only a straightforward distance-based scaling of the original RPY terms, and allows a seamless transition to be made between BD simulations that neglect HIs entirely and simulations that include HIs at the full RPY level of theory.

## Introduction

Molecular simulation studies that employ coarse-grained (CG) structural models of biological macromolecules often make use of implicit solvent methods (e.g. [1,2]). These methods seek to accelerate simulations by eliminating the computational effort that would otherwise be spent on moving solvent molecules that, within the context of CG modeling, are usually of little interest. It is widely recognized that implicit solvent methods must attempt to account for the now-missing solvent’s effects on electrostatic and hydrophobic interactions. A simple approach for dealing with electrostatic interactions within large-scale CG simulations, for example, is to use the Debye-Hückel approximation which models the solvent as a continuum dielectric and which implicitly accounts for the effects of dissolved monovalent ions [3]. A less well appreciated drawback of implicit solvent simulations [4-8] is that they can lead to drastically underestimated diffusional properties of macromolecules. This can severely limit the extent to which they can be used to model macromolecular assembly processes since realistic simulations of such processes require that the relative diffusive properties of monomeric, oligomeric and polymeric species be modeled correctly.

While implicit solvent methods have difficulty modeling the relative diffusive properties of macromolecules correctly, the technique of Brownian dynamics (BD) is unusual in that it can be modified easily (albeit expensively) to solve the problem. The route to this was devised by Ermak and McCammon [9] in work that formulated a rigorous approach for including hydrodynamic interactions (HIs) in BD simulations. HIs are non-energetic terms that describe the solvent-mediated effects that cause nearby solutes to move in a correlated way; importantly, they are long-ranged interactions, decaying with the reciprocal of the distance between solutes. Several studies have shown that including HIs in BD simulations allows the diffusional properties of proteins [4-6] and RNAs [7-8] to be realistically simulated: not only does inclusion of HIs in BD simulations allow the translational (D_trans_) and rotational diffusion coefficients (D_rot_) of globular proteins to be reproduced, for example, it also enables the ∼40% decrease in D_trans_ values that accompany protein unfolding to be reproduced [4]. But the inclusion of HIs in a conventional implementation of the Ermak-McCammon algorithm becomes very expensive as systems increase in size, with the computational cost of large-scale BD-HI simulations often exceeding those of corresponding explicit solvent simulations that they are intended to replace. The exorbitant cost of large-scale BD-HI simulations is probably the major reason they are not more widely used within the biophysics community.

One major contributor to the cost of the BD-HI method developed by Ermak & McCammon [9] is the fact that it allows only an “all or nothing” approach to the modeling of HIs: either one can choose to neglect HIs entirely – in which case, one is performing simulations in the so-called “free draining” regime – or one must include *all* HIs, even those that describe interactions between solute pairs that are separated by enormous distances. Obviously, for extremely distant HIs there are likely to be opportunities to mitigate costs by grouping together solutes that are near to each other and describing their effects on very distant solutes in an approximate manner: this approach lies at the heart of sophisticated fast multipole [10] and treecode [11] methods that have been reported but that have yet to be used for production-level BD-HI simulations.

Here I consider an alternative approach, which is to seek to limit the costs of BD-HI simulations by truncating HIs beyond a specified cutoff distance. Distance-based cutoffs are a common feature of MD simulation codes where they are used to truncate van der Waals interactions beyond a distance at which their contributions are negligible. I am unaware, however, of any reported method that allows a corresponding distance-based cutoff to be used to truncate HIs in BD-HI simulations. The present work shows how a distance-based cutoff for HIs can be implemented: the Methods section first illustrates why the inclusion of a cutoff for HIs is potentially problematic, before showing how the problem can be circumvented in a practical implementation, while the Results section shows how the imposition of a HI cutoff in BD-HI simulations affects the static and dynamic properties of a model protein.

## Methods

### Why a simple distance-based cutoff causes problems in BD-HI simulations

The Ermak-McCammon algorithm [9] requires that the diffusion tensor used to describe HIs has the property of being positive definite, i.e. all of its eigenvalues must be positive. If this is not the case, then it becomes impossible to obtain the correlated random displacements that are needed for a BD-HI simulation to produce correct behavior. To show how the imposition of a simple distance-based cutoff can cause the diffusion tensor to become non-positive definite, consider the system of three particles shown in Figure 1A. A simple possible diffusion tensor for such an arrangement of the particles is indicated by 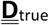. The entries in 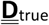 describe the strengths of the inter-particle HIs and reflect the closeness of the particles in space: particles 1 and 2 are close to each other in space, as are particles 2 and 3, so the entries describing their HIs are large in magnitude; particles 1 and 3, on the other hand, are more distant from each other, so the entry describing their HI is small in magnitude. The Cholesky decomposition of this diffusion tensor – obtained using a standard numerical algorithm – is indicated by L; it is triangular matrix that is commonly used to obtain the correlated random displacements needed by the Ermak-McCammon algorithm.

**Figure 1.**
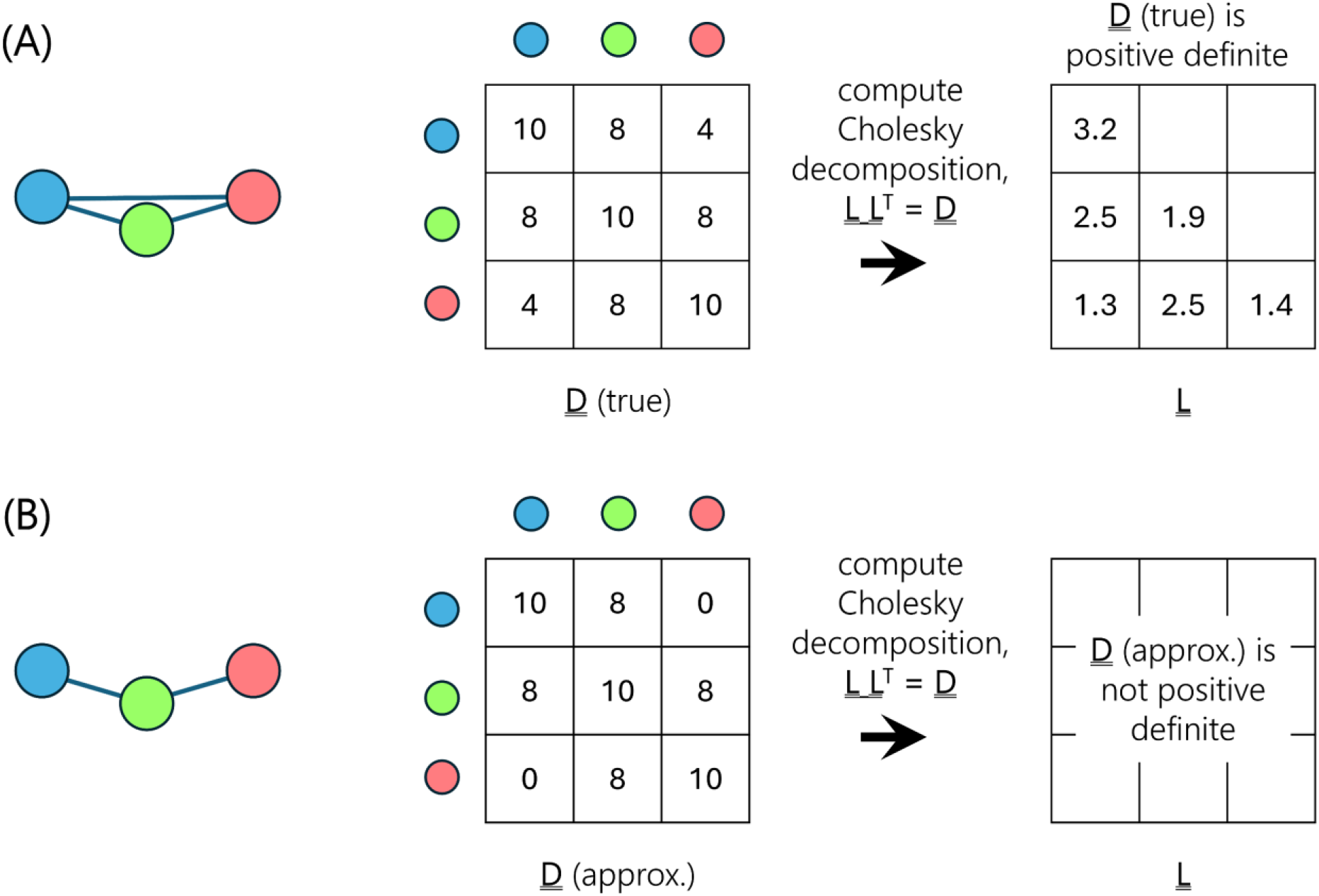
Illustration of the problem with implementing a naive distance-based cutoff. (A) An example three-particle system with its associated (idealized) diffusion tensor, 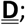 the magnitudes of the entries reflect the strength of the HIs acting between the particles. The Cholesky decomposition for this 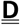 is shown at right. (B) Same as (A) but with the long-range HI between particles 1 and 3 set to zero to illustrate the effect of applying a simple distance-based cutoff. The Cholesky decomposition for this 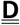 is not positive definite and therefore does not contain purely real components.

In Figure 1B we show what happens if we naïvely truncate long-range HIs with a distance-based cutoff. We consider a situation in which the cutoff distance is set such that the HI between particles 1 and 3 lies beyond the cutoff and is therefore set to zero, while the two remaining interactions (1-2 and 2-3) are within the cutoff and are left unchanged. The problem with this approach becomes apparent when we try to compute the Cholesky decomposition, 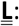 the numerical algorithm fails to produce a result since 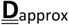, is now non-positive definite. This in turn prevents the computation of the correlated random displacements that would be used to propagate motion of the particles via the Ermak-McCammon algorithm.

### A solution to the problem

We illustrate the solution to the above problem by considering the somewhat more complicated system of six particles shown in Figure 2. A possible simplified representation of the “true” diffusion tensor for this system is shown in Figure 2A; again, the magnitudes of the entries in the tensor reflect the closeness of the particles in space. We now show how to implement a distance-based cutoff in such a way that the resulting approximate diffusion tensor is guaranteed to remain positive definite. We proceed by first considering some possible ways to partition the complete six-particle system into smaller sub-systems in such a way that: (a) all HIs involving particle pairs that reside *within the same sub-system* are computed in the usual way with the RPY level of theory, and (b) all HIs involving particle pairs that reside *within different sub-systems* are neglected entirely, i.e. set to zero. The simplest way to carry out this kind of partitioning is to place the particles onto a cubic grid that is dimensioned such that the distance from one corner of each grid cell to the furthest corner is equal to the desired HI cutoff distance (for a cutoff distance of 20 Å, for example, the grid cells should have sides of length 11.547 Å). The partitioning of the particles into the grid cells, and by extension their HI sub-systems, will obviously depend on where exactly the origin of the grid is placed. Figure 2B shows five example partitionings of the system, together with the resulting realizations of the diffusion tensor (see bottom row of Figure 2B). Importantly, all five of the diffusion tensors shown are guaranteed to be positive definite since they each can be considered as collections of smaller diffusion tensors that either: (a) contain sub-sets of particles for which all of the possible HIs are computed, or that (b) contain only a single particle. This is the case even for the fourth possible realization shown which has particles 1, 2, and 5 forming a single sub-system and particles 3 and 4 forming a separate sub-system: a simple reordering of the elements of the diffusion tensor will make clear that this tensor remains positive definite.

**Figure 2.**
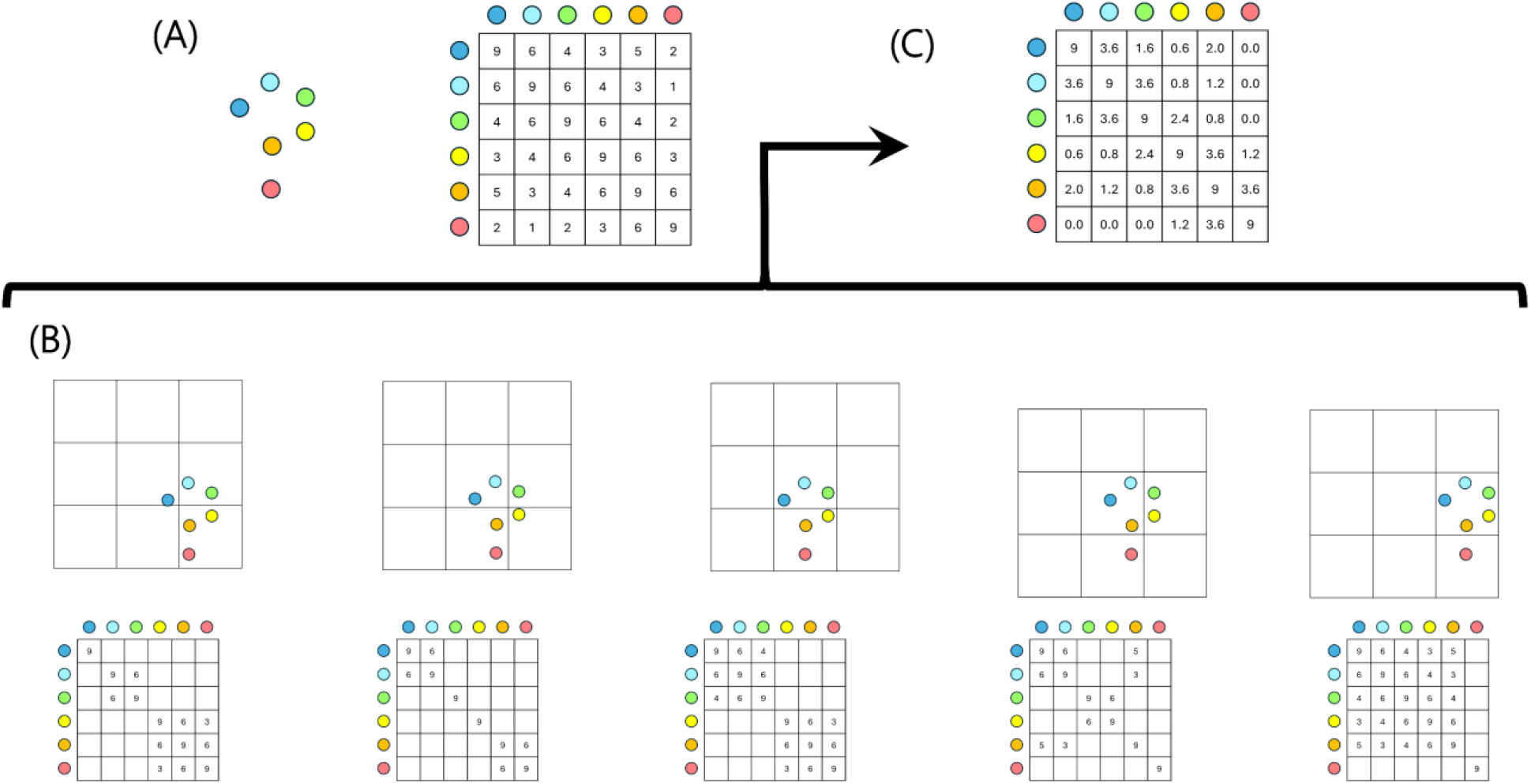
Example illustration of how a distance-based cutoff can be implemented. (A) An example six-particle system with its associated (idealized) diffusion tensor, 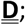 the magnitudes of the entries reflect the strength of the HIs acting between the particles. (B) (top row) five possible partitionings of the system obtained by placing particles onto a cubic grid; (bottom row) the corresponding diffusion tensors obtained when HIs are included only between particles that are part of the same sub-system. (C) approximate diffusion tensor obtained from arithmetic averaging of the five diffusion tensors shown in (B); this diffusion tensor truncates long-range HIs while remaining positive definite.

Crucially, a matrix that is the arithmetic average of matrices that are *all* known to be positive definite is guaranteed also to be positive definite, so it is possible to simply average all of the entries in the five different diffusion tensors to obtain a final approximate diffusion tensor shown in Figure 2C. The resulting approximate tensor has the longer-range HIs correctly set to zero beyond the cutoff distance and is guaranteed to remain positive definite; moreover, it is clear from examination of the entries that the relative strengths of the HIs in the approximate diffusion tensor are qualitatively similar to their strengths in the “true” diffusion tensor shown in Figure 2A. The latter result is a consequence of the fact that when the system is partitioned into sub-systems by mapping to a cubic grid, pairs of particles that are close to each other in space are more likely to partition to the same sub-system and therefore have their HIs retained. In contrast, pairs of particles that are more distant from each other are more likely to partition to different sub-systems and therefore have their HIs omitted.

### Numerical determination of cutoff function derived numerically using cubic cells

A more complete treatment of the system shown in Figure 2 would require that all possible origins for the cubic grid be considered, and that all possible orientations of the system relative to the cubic grid also be considered. The approximate diffusion tensor obtained by averaging all possible partitionings in this way would be identical to the true diffusion tensor but with the entries for each pair of particles scaled by the mean probability that they partition into the same sub-system. In what follows, I will term this an acceptance probability, P_acc_, and I will write it as a function of the ratio of the actual distance between the particles, l, to the cutoff distance l_cut_, i.e. as P_acc_(l/l_cut_). In effect, P_acc_ reflects the probability that a line of relative length, l/l_cut_, that has one end placed at a random location within a grid cell (whose cross-diagonal distance is l_cut_), has its other end also contained within the same grid cell. While there may be an analytical solution to the problem of calculating P_acc_ as a function of l/l_cut_, I have chosen here to evaluate it numerically, using for each possible value of the separation distance, 10 million randomized orientations of each possible line length.

### Implementation within Brownian dynamics simulations

With P_acc_ as a function of l/l_cut_ determined numerically (see Results), a distance-based cutoff can be applied to BD-HI simulations that use the RPY diffusion tensor straightforwardly. Relative to the conventional BD-HI simulation approach in which no cutoff is applied, only two changes are required. First, for each pair of particles whose inter-particle distance lies within the HI cutoff distance, the terms of the Rotne-Prager-Yamakawa (RPY) diffusion tensor [12,13] are computed as usual and multiplied by the value of P_acc_ that corresponds to the separation distance of the two particles. Second, since scaling the RPY terms in this way causes the diffusion tensor to have a non-zero divergence term, care must be taken to calculate it and include it in the position-update algorithm as outlined by Ermak and McCammon [9]; a similar issue was encountered in our recent work implementing an orientationally averaged version of the RPY tensor [14]. In the present case, the divergence term can be calculated using the product rule for differentiation: the divergence of the diffusion tensor is equal to the RPY terms multiplied by the slope of P_acc_ versus distance, which we again determine numerically.

### Brownian dynamics simulations of a model protein

All simulations reported here were carried out with the uiowa_bd code (https://github.com/Elcock-Lab/uiowa_bd) with minor modifications made to allow for the use of a distance-based cutoff. BD-HI simulations were tested on a single protein, the 149-residue Semlike Forest Virus Protein (SFVP), modeled at a resolution of one “bead” per residue. Simulations were carried out with the parameters used in our previous study that tested the use of the orientationally averaged form of the RPY tensor for modeling translational and rotational motions of proteins [14]. Briefly, bonds between consecutive residues were maintained at their equilibrium lengths using harmonic potentials with a force constant of 20 kcal/mol/Å^2^; angles involving three consecutive residues were maintained using harmonic potentials with a force constant of 10 kcal/mol/rad^2^; dihedral angles involving four consecutive residues were treated with cosine potential functions with periodicities of 1 and 3, and with energy barriers of 0.5 and 0.25 kcal/mol respectively. In simulations of the protein in its folded state, all nonbonded residue pairs for which at least one of their atom pairs was within 5.5 Å in the native state were assigned a favorable Lennard-Jones potential with a well-depth of 1 kcal/mol; all other residue pairs were assigned purely repulsive 1/r^12^ potential function with a distance parameter (σ value) of 4 Å and an energy parameter (ε value) of 0.1 kcal/mol. All BD-HI simulations were performed with the Ermak-McCammon algorithm with a timestep of 25 fs for 10 µs, with snapshots of the protein saved at intervals of 1 ns. The diffusion tensor, together with its Cholesky decomposition, and the associated divergence term, was updated every 100 simulation steps.

## Results

### Acceptance probability versus distance

As described in Methods, a key step in developing a distance-based cutoff for BD-HI simulations is to determine an acceptance probability function, P_acc_, that quantifies the probability that two particles will be partitioned into the same hydrodynamic sub-system. Here we express P_acc_ as a function of the ratio of the inter-particle distance to the cutoff distance (l/l_cut_), and we derive its distance dependence numerically by partitioning the system of interest into cubic grid cells (see Methods). The resulting P_acc_ function is shown in Figure 3. The function behaves broadly as expected. At one end of the scale, P_acc_ approaches 1 as l/l_cut_ tends to 0, thereby reflecting the fact that pairs of particles that are extremely close to each other are highly likely to partition into the same hydrodynamic sub-system: for such particle pairs, the effective strength of the HI applied in the proposed cutoff scheme will be very close to that obtained without any cutoff. At the other end of the scale, P_acc_ approaches 0 as l/l_cut_ tends to 1, i.e. as the inter-particle separation distance approaches the cutoff distance; for values of l/l_cut_ above 1, P_acc_ is, by definition, 0. Notably, P_acc_ is close to 0 even for inter-particle distances that are considerably shorter than the cutoff distance.

**Figure 3.**
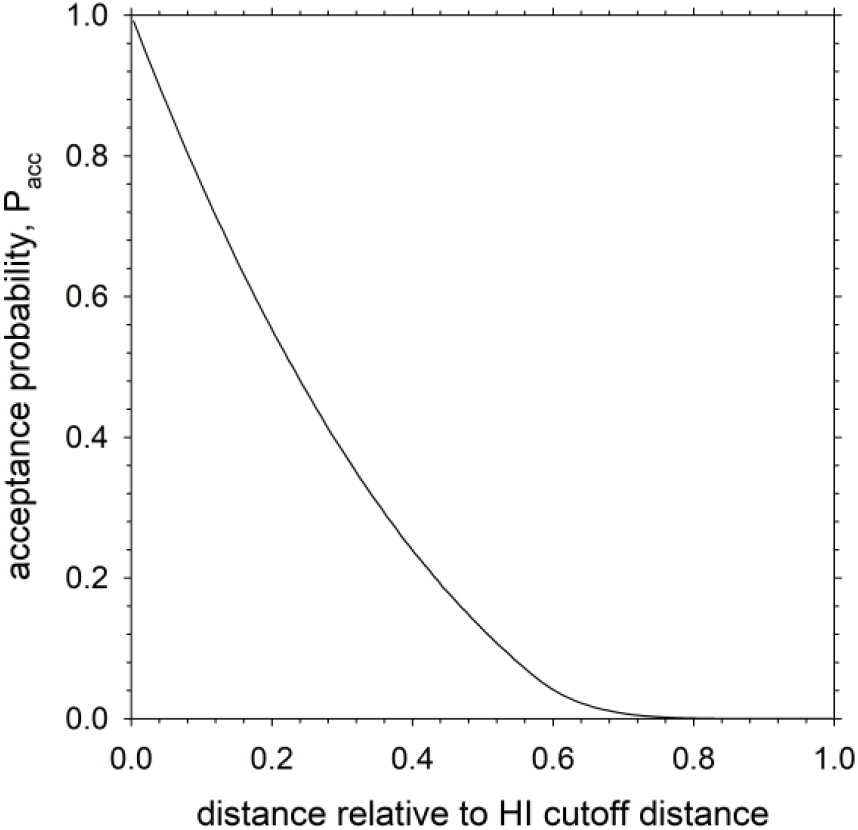
Acceptance probability, P_acc_, plotted as a function of the reduced distance l/l_cut_.

### Static properties of a model protein are unaffected by the inclusion of an HI cutoff

A critical test of any approximate BD-HI algorithm is that it produces correct behavior. In the context of highly flexible systems such as unfolded or disordered proteins, the simplest way to demonstrate this is to examine static properties such as the radius of gyration and show that the value obtained from BD-HI simulations is independent of the parameters of interest. Figure 4A shows that this is indeed the case for the model protein SFVP, a 149-residue protein, that we have studied previously [14]. The figure plots the mean radius of gyration of the protein in both its folded and unfolded states as obtained from 10 µs BD-HI simulations that employ different values of the cutoff distance for HIs. The left-most datapoint labeled “no HI” corresponds to a cutoff distance of 0 Å and refers, in effect, therefore to a BD-HI simulation performed in the absence of any HIs, i.e. within the free-draining approximation; the right-most datapoint labeled “no cutoff” indicates a BD-HI simulation performed in the presence of all HIs, even those that act between particles that are widely separated by distance, and with all such HIs unscaled by P_acc_.

**Figure 4.**
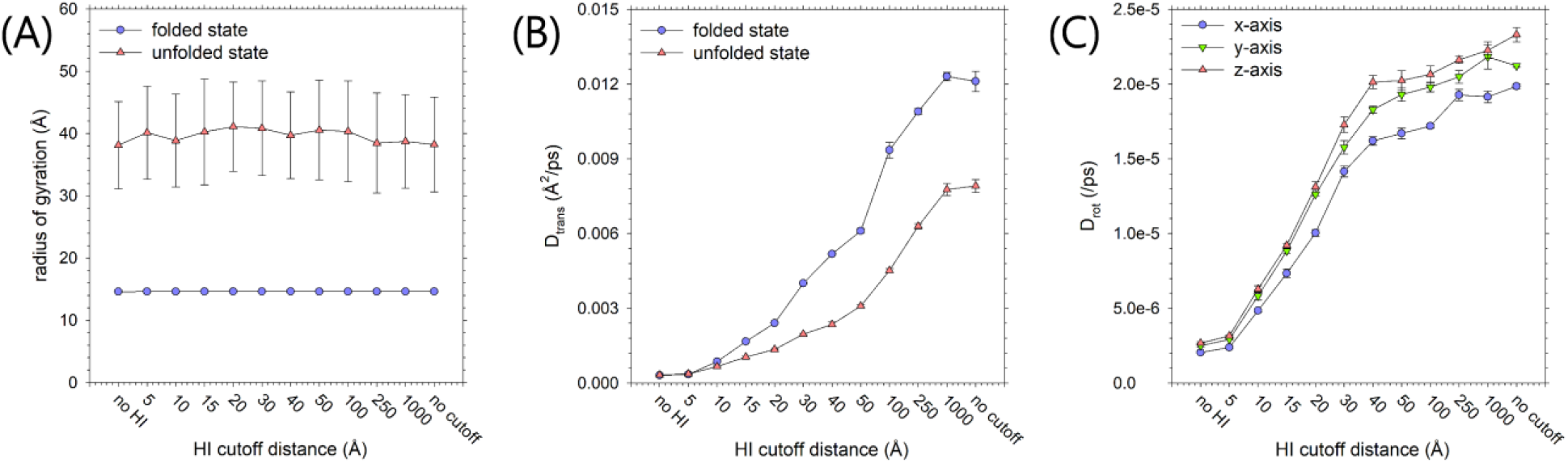
Static and dynamic properties obtained from BD-HI simulations with a distance-based cutoff. (A) Mean radius of gyration obtained from 10 µs BD-HI simulations of the model protein SFVP in its folded (blue) and unfolded (red) states plotted as a function of the cutoff distance; error bars represent the standard deviation of 10,000 sampled conformations. (B) Mean translational diffusion coefficients of SFVP in its folded (blue) and unfolded (red) states plotted as a function of the cutoff distance; error bars represent the standard deviation of the mean values obtained from three contiguous blocks of data (C) Same as (B) but showing mean rotational diffusion coefficients of the three principal axes of SFVP in its folded state.

Importantly, regardless of the applied cutoff distance, the results shown in Figure 4A indicate that the resulting radius of gyration is unchanged and that this is true regardless of whether the protein is modeled in its folded (blue) or unfolded (red) states.

### Diffusional properties of a model protein as a function of HI cutoff distance

We know from previous work that the inclusion of HIs modeled at the RPY level of theory is needed for BD simulations to accurately simulate the translational and rotational diffusion of CG macromolecules and that the omission of HIs results in both translational and rotational diffusion being hugely underestimated. We should expect, therefore, that replacing the usual RPY diffusion tensor with one calculated with a distance-based cutoff will negatively impact results, with behavior getting progressively worse as the cutoff distance is decreased. We show that this expectation is fulfilled in the remaining panels of Figure 4. Figure 4B plots the translational diffusion coefficient, D_trans_, versus cutoff distance for SFVP in its folded (blue) and unfolded (red) states. At low values of the cutoff distance the two diffusion coefficients are effectively equal: this reflects the known result that BD simulations that omit HIs are incapable of reproducing the difference in translational diffusion of folded and unfolded states of proteins [4]. For the folded state of the protein, the D_trans_ value obtained when all HIs are omitted is ∼40 times lower than the value obtained when all HIs are included and unscaled. As the cutoff distance increases, the D_trans_ values of the folded and unfolded states gradually increase and diverge as progressively longer-range HIs are included in the simulations. For intermediate values of the cutoff distances, the D_trans_ value of the folded state appears to increase rather more than that of the unfolded state; this presumably reflects the fact that it has a greater number of short-range HIs than the unfolded state. For both the folded and unfolded states, a cutoff distance of 50 Å is required for the D_trans_ values to approach 50% of its value obtained when all HIs are included and unscaled.

A similar picture emerges from analysis of the rotational diffusion coefficients of SFVP, which are plotted for each of the three principal axes of the protein in its folded state in Figure 4C. For each of the three axes, the D_rot_ values obtained when all HIs are omitted are ∼9-10 times lower than the values obtained when all HIs are included and unscaled: the finding that rotational diffusion is somewhat less affected by the omission of HIs than translational diffusion is consistent with previous work [4]. As expected, increasing the cutoff distance results in a gradual increase for all three D_rot_ values, but as was also the case with translational diffusion (Figure 4B), a cutoff distance of ∼50 Å is needed before they reach 50 % of the values obtained when all HIs are included and unscaled.

### Minimum and maximum eigenvalues of the diffusion tensor as a function of cutoff distance

In all of the test simulations described here, the correlated random displacements required by the Ermak-McCammon algorithm were obtained by first calculating the Cholesky decomposition of the diffusion tensor at regular intervals. In practical applications, a HI cutoff approach is expected to be accompanied by the use of iterative methods for calculating correlated random displacements since these can be applied to much larger systems than the Cholesky decomposition-based approach. Two commonly used iterative methods are the Chebyshev polynomial-based method first proposed by Fixman [15,16], and the Krylov subspace-based method developed by the Chow and Skolnick groups [17]. For the Chebyshev polynomial method, the number of iterations required for convergence of the correlated random displacements has been shown to increase as the so-called “condition number” of the diffusion tensor increases [16]. An important point to explore, therefore, is how the condition number of the diffusion tensor is affected by the imposition of a distance-based cutoff to HIs. This is addressed in Figure 5, which plots the condition number of SFVP’s folded state diffusion tensor as a function of cutoff distance. As with the other plots included here, the condition number gradually increases as the cutoff distance increases in an effectively monotonic manner; this straightforward dependence suggests that the imposition of a cutoff will accelerate the rate at which correlated random displacements are obtained in a predictable manner.

**Figure 5.**
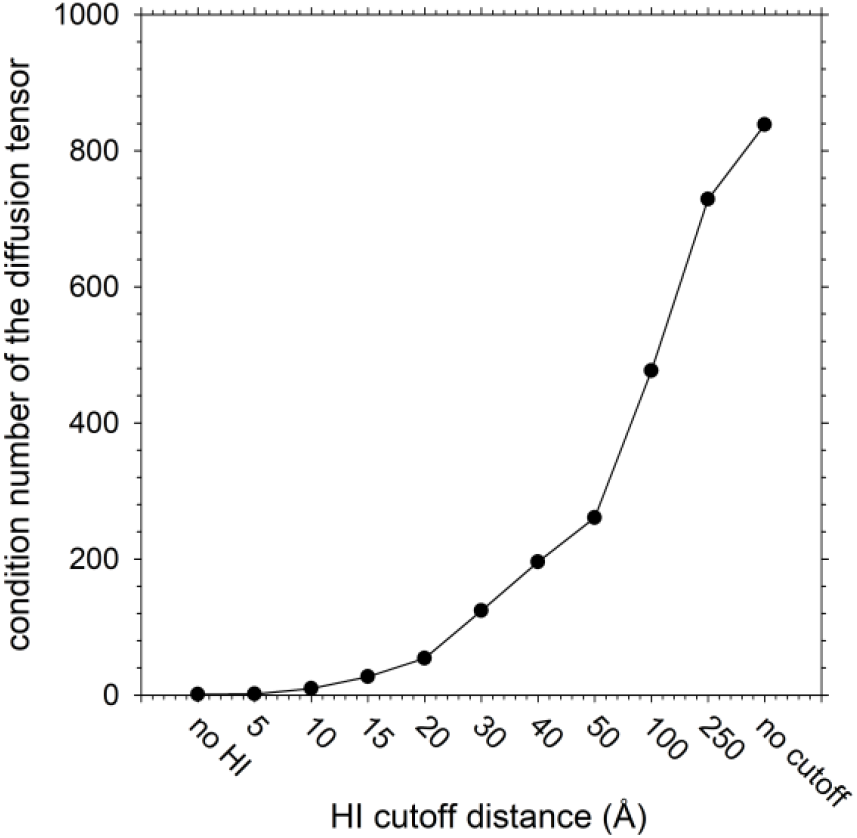
Condition number of SFVP’s folded state diffusion tensor plotted as a function of the cutoff distance. The condition number is computed as the ratio of the largest and smallest eigenvalues of the diffusion tensor.

## Discussion

This work has described a straightforward way to implement a distance-based cutoff to HIs in BD-HI simulations. The simplest possible approach, which is to set elements of the diffusion tensor to zero for those particle pairs that are further apart than the cutoff distance, has a clear potential to cause the diffusion tensor to lose its critical characteristic of being positive definite (see Methods). But this problem can be avoided using the approach outlined in Methods, in which the system of interest is conceptually divided up into smaller sub-systems that are each modeled in such a way that guarantees that the diffusion tensor remains positive definite. It is to be noted that this approach adheres to the “all or nothing” approach to modeling HIs that was outlined in the Introduction: all of the possible HIs within a given sub-system are included. Generalizing this idea into a practical implementation requires only the introduction of an acceptance probability, P_acc_, that progressively scales down the RPY tensor terms as the inter-particle separation distance approaches the cutoff distance.

In application to a simple protein system the inclusion of a cutoff to HIs has been shown to have both advantages and disadvantages. On the negative side, the imposition of a progressively shorter cutoff distance clearly decreases the computed D_trans_ and D_rot_ values, thereby making them deviate further and further from the realistic values obtained when no HI cutoff is applied (Figures 4B and 4C). At a qualitative level, this trend is to be expected given that in the limit of a zero cutoff distance the approach outlined here yields a simulation model in which all HIs are omitted. But at a quantitative level, the large decreases in the computed D_trans_ and D_rot_ values are likely to be due in part to the shape of the P_acc_ function (Figure 3), which suppresses HIs substantially even at inter-particle distances that are considerably shorter than the cutoff distance.

On the positive side, the imposition of a progressively shorter cutoff distance has the potential to significantly decrease the computational expense associated with performing BD-HI simulations on large-scale systems. Decreases in costs are expected for three reasons. First, since the imposition of a cutoff will zero out a large number of elements of the diffusion tensor, 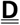, the cost of multiplying 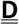 by the force vector 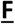 – an operation that must be carried out at each time step of the simulation [9] – will, at least in principle, decrease substantially since the dense matrix-vector product required when no cutoff is applied will be replaced a sparse-matrix vector product. Obviously, the expected speedup will depend on how sparse 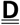 becomes, and this in turn will be determined by the choice of cutoff distance. Second, by the same argument, the imposition of a cutoff will decrease the cost of computing correlated random displacements using iterative methods such as the Chebyshev polynomial [15,16] or Krylov subspace [17] methods: both methods involve the use of repeated matrix-vector product calculations in which the matrix is a scaled version of 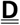. Third, the cost of computing correlated random displacements is likely to be further reduced by the fact that the condition number of 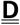 decreases substantially as the cutoff distance decreases (Figure 5): this means that fewer iterations will be required to obtain converged correlated random displacements.

Since there are significant advantages and disadvantages associated with the distance-based cutoff method proposed here, its utility in practical situations will depend on whether the decreased costs associated with imposing a cutoff outweigh the poorer description of translational and rotational diffusion that is likely to result. Although the answer to that question will be a function of the system under study, the method developed here provides a route to seamlessly incorporating HIs at all possible length-scales, ranging from a length-scale at which they are omitted entirely to one at which they are included at full strength.

## Acknowledgments

This research was supported by the University of Iowa and by a NIH grant (R35 GM122466) to AHE.

## Author Contributions

Adrian H. Elcock: Conceptualization, Methodology, Software, Investigation, Writing = Original Draft, Supervision, Project Administration, Funding acquisition.

## Data availability

All simulations were performed with a modified version of the uiowa_bd code [14] that is hosted at the following GitHub repository: https://github.com/Elcock-Lab/uiowa_bd.

